# Practical bioinstrumentation developments for AC magnetic field-mediated magnetic nanoparticle heating applications

**DOI:** 10.1101/328211

**Authors:** Mahendran Subramanian, Arkadiusz Miaskowski, Ajit K. Mahapatro, Ondrej Hovorka, Jon Dobson

## Abstract

Heat dissipation during magnetization reversal processes in magnetic nanoparticles (MNP), upon exposure to alternating magnetic fields (AMF), has been extensively studied in relation to applications in magnetic fluid hyperthermia (MFH). This current paper demonstrates the design, fabrication, and evaluation of an efficient instrument, operating on this principle, for use as (i) a non-contact, *in vitro*, real-time temperature monitor; (ii) a drug release analysis system (DRAS); (iii) a high flux density module for AMF-mediated MNP studies; and (iv) an *in vivo* coil setup for real-time, whole body thermal imaging. The proposed DRAS is demonstrated by an AMF-mediated drug release proof-of-principle experiment. Also, the technique described facilitates non-contact temperature measurements of specific absorption rate (SAR) as accurately as temperature measurements using a probe in contact with the sample. Numerical calculations estimating the absolute and root mean squared flux densities, and other MNP – AMF studies suggest that the proposed stacked planar coil module could be employed for calorimetry. Even though the proposed *in vivo* coil setup could be used for real-time, whole body thermal imaging (within the limitations due to issues of penetration depth), further design effort is required in order to enhance the energy transfer efficiency.

## Introduction

The current aspiration of magnetically targeted/triggered drug and biomolecule delivery is to minimize the quantities of drugs used in therapies and to more effectively deliver them to the therapeutic target. The demands of modern instrumentation mean that high precision tools are needed in order to overcome issues when developing and standardising experimental methodologies [Kozissnik, B et al., 2013]. This influences the ability of future technologies to meet the demand for the accurate detection of electrical, magnetic, optical, and thermal signals using sensitive, high-precision tools. In the past decade, alternating magnetic field (AMF) mediated magnetic nanoparticle (MNP) heating has attracted a great deal of attention due to its potential in biomedical applications.

Non-contact, *in vitro* temperature measurement techniques, such as infra-red thermography and IR sensors, has been difficult due to the requirement for direct optical exposure to the sample. Scientists and engineers from multidisciplinary research areas have developed instruments which use very specific algorithms [Khan, M. A et al., 1991]. These needs to be standardized so that well-evaluated and easy to use tools, compatible with biomedical applications, can be designed. With the currently available instruments, the temperature sensing used for the magnetic fluid calorimetry is recorded by utilizing a thermocouple, an infra-red sensor, infra-red thermography, or fibre optics sensors [Skumiel, A et al., 2016]. However, each of these techniques has its own advantages and disadvantages. Thermocouples are made up of metals (copper and constantine) and when exposed to AMF, metals generate heat due to Eddy currents, produce radio-frequency (RF) interference, and develop corrosion after repeated use when they are immersed within the colloid. Contamination of samples is also possible when using the same thermocouple for repeated calorimetric measurements. This means that metal thermocouple probes are not really usable in the presence of AMF [Rose, L. C. et al 2016; Elkhova, T. M. et al 2014; Rahn, H et al., 2013]. Infra-red thermography and infra-red sensor modules require direct optical exposure to the *in vitro* sample [Skumiel, A et al., 2016; Rodrigues, H. F. et al 2013]. This involves heat loss from the sample and carries the risk of sample contamination. Similarly, a fibre optic sensor is not prone to electromagnetic interference but, as mentioned above, does have to have direct contact with the sample. In addition, these are expensive, delicate (silicone tube), and the probability of cross contamination is high. This is one of the reasons for the limited number of experiments using real-time temperature measurements in magnetic fluid hyperthermia research involving cell cultures; such experiments require a contamination-free environment [Rodríguez, H. L. et al 2009; Subramanian, M et al., 2016; Subramanian, M. et al 2016]. MRI mediated non-contact temperature measurement is plausible, but, customisation, accessibility, ease of use and cost involved require consideration [Fossheim SL et al., 2000].

AMF-mediated drug delivery is popular because of the fact that it is non-invasive and remotely triggered. Controlled drug delivery systems have been a broad research interest for a number of years. The stimuli used here is the heat dissipated from surface functionalised magnetic nanoparticles exposed to AMF [Satarkar, N. S. & Hilt, J. Z 2008; Teleki, A et al., 2016; Ding, X. et al 2016]. There are a few crude setups for performing such experiments using commercially available AMF systems [Brulé, S. et al 2011; Hervault, A. et al. 2016], and the research community has discussed the prospects for the development of nanoparticle-based drug delivery testing apparatus [D’Souza, S et al., 2014]. However, this requires certain modifications to be made to the United States Pharmacopeia (USP) type drug testing setups generally used in the pharmaceutical industry. Such modifications would have to take the following into account: nanoscale level fillers/binders, release patterns, the stimuli involved, the AMF generator, stirring mediated convection loss, precise sample positioning, the requirement for a water jacket, and the sampling method. The instrument development considered here looks at USP and flow-cell type apparatus in relation to the designing and fabrication of AMF-mediated drug delivery testing setups which facilitate precise physical and chemical measurements.

Biomedical application research within this topic has advanced significantly over the past decade, however robust and well evaluated tools that will allow researchers to replicate experiments is a must. Herein, we provide insights into development of practical modules for commercialization. This paper presents our recent innovative advances in bioinstrumentation research and demonstrates four different elementary but effective tools that could benefit researchers working in fields related to AMF-mediated MNP heating. These include; (1) non-contact *in vitro* temperature measurement; (2) drug release analysis setups; (3) the utilization of stacked planar coils with increased flux density at low current input for calorimetric measurements; and (4) *in vivo* coil setups for real time full body thermal imaging.

Preliminary testing of our non-contact temperature measuring technique for the evaluation of specific absorption rate (SAR) estimated 8.88 ± 0.24 W/g for a test MNP containing 10 mg / ml aqueous suspension of dimercaptosuccinic acid stabilised magnetic nanoparticles. These results were in good agreement with the value of 8.83 ± 0.15 W/g as yielded by a temperature measurement probe which was placed in contact with the MNP. Likewise, we have designed, fabricated, and evaluated an AMF mediated drug release testing setup; a 2-layer stack of planar coils that provides increased flux density for calorimetric experiments (with no harmonics at low current input). This module generates high flux density with low coil current, uses high frequencies, and provides uniform and homogeneous incident flux density to expose onto the calorimetric sample. This stacked planar coil system was evaluated, in combination with an associated high voltage variable capacitance module, for its ability to produce electromagnetic frequencies between 50 kHz to 250 kHz. Additionally, we built an *in vivo* coil setup embedded within a small animal bed, with water jacket and a viewing window, for real time - whole body thermal imaging.

### Methods and materials

CAD was used to design the systems and components were fabricated using precision engineering and 3D printing. Physical measurements of MNP heating under RF exposure and numerical model based calculations were performed to evaluate the effectiveness of the proposed techniques and design.

### Non-contact temperature measurement

The Live cell – Alternating Magnetic Field^TM^ (LCAMF) module connected to a magneTherm^TM^ system enables RF exposure to an MNP sample in a petri dish (Lumox^®^ 35 mm, Sarstedt, Numbrecht, Germany), with or without cells [Subramanian, M 2016]. An ultra-thin polystyrene (or other polymer) film in a cell culture petri dish is formed as a base, and used as a window for gas exchange and non-contact temperature measurements. The fibre optic temperature sensor (OTG-M360-10-62ST-1.5PTFE-XN-10GT-M2; OPSENS, Quebec, Canada) measurements were undertaken by attaching the sensor tip to the bottom of the petri dish; this allowed the recording of the temperature of the underside of the film. In the case of the mid-infrared temperature sensor (CSmicro 2W Mid-Infrared (MIR), Optris, Berlin, Germany), with a working range of 8-14 μm, the sensor head was situated on the underside of the film.

### Computer aided design (CAD) and computer aided Machinery

Drug release analysis system and coil forms were designed using CAD. All CAD work was done using PTC Creo v3.0 (Staffordshire, UK) and the 3D printing was depicted using Flashforge-Creator pro (London, UK).

### Stacked, curved planar coil fabrication

Hollow water-cooled coils were fabricated using 18 standard wire gauge (SWG) copper wire (Maplin, Stoke on Trent, UK) and silicon tube (Saint-gobain, Coventry, UK) with dimensions 4 mm ID × 6 mm OD. A coil form (bobbin) was designed and precision engineered for use in winding and fixing the geometry of the coil.

### Microprocessor programming

The interfacing software was programmed in C++ on an off-line development system. The software was run on an Arduino Uno microprocessor board driving a motor shield v1.0 (Farnell, Angus, UK). The Arduino IDE platform was used to upload the firmware onto the microprocessor.

## Experimental and numerical evaluation

### Calorimetric experiment

20 mg/ml aqueous suspensions of 50 nm sized maghemite (Sigma Aldrich, Dorset, UK) and 10 mg/ml aqueous suspensions of dimercaptosuccinic acid (DMSA) stabilized 10.3 nm sized magnetite (Hypermag A, nanoTherics, Staffordshire, UK) were used for the calorimetric and drug release testing measurements. Identical containers were used in all the calorimetric experiments. A Lumox^®^ 35 mm petri dish (Sarstedt, Numbrecht, Germany) was used for the *in vitro* temperature measurements and a 3 ml Pur-A-Lyser dialysis tube (Sigma Aldrich, Dorset, UK) was used for testing the drug release. MNPs were subjected to vortex and ultra-sonicate prior to AMF exposure. A Pico M^™^ with OTG-MPK 5 optical sensor system (Opsens, Quebec, Canada) and an Osensa fibre optic temperature sensor (Burnaby, Canada) were used for recording real-time temperature measurements. An alternating current (AC) magnetic field – magneTherm system (nanoTherics, Staffordshire, UK) was used as a source of exposure. The specific loss power (W/g of particles) was calculated using the following equation:

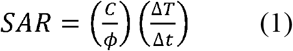

where C is the specific heat capacity of the MNP in J/Kml, □ is the concentration of MNP in mg/mL, and Δ*T/*Δ*t* is the rate of change of temperature over time. An appropriate region of the graph was utilized, using the corrected slope method, for calculations [Wildeboer, R. R et al., 2014]. The baseline temperature was subtracted from the results, and the linear loss parameter was estimated from the temperature difference over time slope – based on fitting interval and number of fits.

### Spectroscopy measurements

Absorption spectroscopy was performed using a Shimadzu UV-Vis spectrophotometer (Milton Keynes, UK); the samples were placed in a quartz crystal cell. Real time UV-Vis absorption spectroscopy data was recorded using a non-metal fibre optic probe (FCB-UV200-1.5-S1, Avantes, Surrey, UK) connected to a halogen light source (Visual and near infra-red range, Avantes, Surrey, UK) and a spectrophotometer module (Visual range, Avantes, Surrey, UK) while performing the AMF mediated drug release experiment. The cancer drug – doxorubicin hydrochloride – was procured from Sigma Aldrich (Dorset, UK).

### Flux density simulation and calculation

The magnetic flux density (B) was simulated. This was done by the Sim4Life (ZMT Zurich MedTech, Zurich) platform. The flux distribution was modelled in relation to an applied low frequency magneto-quasi static (LF M-QS) algorithm utilizing the finite element method (FEM) model. Here, the coil wire was treated as a perfect electric conductor (PEC) and the parameters defining the coil geometry used in the numerical modelling included: the outer diameter of the tube (D_t_), the inner diameter of the tube (d_t_), the use of 18 standard wire gauge copper wire (g_cu_), the number of turns (N), the number of layers (l), the distance between the turns (S_N_), the distance between layers (S_1_), the inner diameter of the coil (d_c_), and the outer diameter of the coil (D_c_). Two coils were modelled i.e. the double stack planer one and the curved planar one.

For the double stacked planar coil with 14 turns each the simulation with D_t_ = 6 mm, d_t_ = 4 mm, g_cu_ = 1.22 mm, N = 14, l = 2, S_N_ = 3.208 mm, S_l_ = 6.78 mm, d_c_ = 17.208 mm, and D_c_ = 132.336 mm.

For the curved planer modelling included: D_t_ = 6 mm, d_t_ = 4 mm, g_cu_ = 1.22 mm, N = 11, l = 1, SN = 4 mm, d_c_ = 5 mm, D_c_ = 60 mm, and C_i_ (internal curvature) = 90 mm. In this case we presented the coil mathematically as an Archimedean spiral then transformed its planar geometry to circular geometry so that it would appear to lie on a plastic cylinder.

## *In vitro* - numerical modelling

### Eddy current effect

When human tissues are exposed to an alternating electromagnetic field, the eddy current effect is observed due to non-zero conductivity (σ) of the tissues, which ultimately leads to their heating [S. Dutz and R. Hergt 2013]. In order to include a volumetric power density (*P*_*eddy*_) produced by an eddy current (*J*_*eddy*_) effect, a magneto-quasi static algorithm was applied [Miaskowski, 2012]. *P*_*eddy*_ can be expressed as:

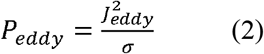

Discussing the eddy or Foucault currents is essential due to their effects on different types of tissues when exposed to a time-varying magnetic field. The eddy current effect can lead to unwanted non-specific heating of healthy tissues, so must be considered in the applicator design.

### Single-domain magnetic power losses

In addition to the power losses due to the eddy current effect, a power dissipation from magnetic nanoparticles (MNPs) should be considered in parallel when dealing with MFH. When single-domain MNPs in the paramagnetic regime are exposed to an AMF with given parameters (*H*_*max*_, *f*), the magnetization lags behind the external field thus specific loss power can be express as

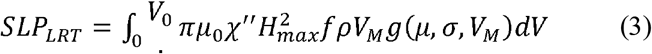

where *ρ* is the density of MNPs *χ″* and is the average out-of-phase component of the susceptibility given by

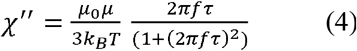

where τ is the Neel-Brown relaxation time *k*_*B*_ is the Boltzmann constant, *T* is the temperature, *μ* is the magnetic moment of the magnetic particle defined as *μ* = *M*_*S*_*V*_*M*_ where *M*_*S*_ stands for the saturation magnetization and *V*_*M*_ stands for the magnetic volume of the nanoparticle 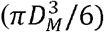. Taking into account polydisperse character of MNPs expressed as the volume weighted distribution *g*(*μ*, *σ*, *V*_*M*_) together with the Zeeman condition one can find a critical diameter/volume above which the MNPs will blocked. In this case specific loss power can be expressed as

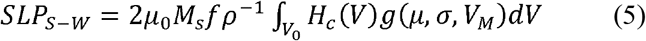

where *H*_*c*_ is the coercively [Miaskowski A et al., 2016; Miaskowski A and Subramanian M., 2017; Miaskowski A et al., 2018].

### Pennes equation

In this case, the heat generation rate, the perfusion and the heat-transfer rate were made linearly dependent on the temperature (*aT* + *b*) to account for the strong temperature dependence due to bio-regulatory processes. Hence, temperature distribution in the breast model was investigated using the modified bio-heat transfer equation formulated by [Pennes, H. H 1948]:

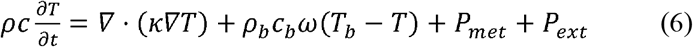

where *ρ* is the tissue density [kg/m], *c* is the tissue specific heat [J/kg/C], *ρ*_*b*_ is the density of blood [kg/m3], *c*_*b*_ is blood specific heat [J/kg/C], *κ* is the tissue thermal conductivity [W/m/C]), *ω* is the perfusion rate [1/s], *T*_*b*_ is the arterial blood temperature [C], *T* is the local temperature [C], *P*_*met*_ is the metabolic heat source [W/m^3^] and *P*_*ext*_ = *ρ*_*MNP*_(*SLP*_*LRT*_ + *SLP*_*S-W*_) + *P*_*eddy*_ is the external heat due to eddy currents and the magnetic nanoparticles power losses [W/m^3^]. The convection boundary condition (Robins type) was used:

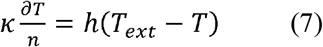

where *h* is the heat transfer coefficient and κ is the thermal conductivity.

### *In vivo* - numerical modelling

The thermal properties of the rat tissue model were calculated for *f* = 107 kHz using 4-Cole-Cole approximation based on IT’IS database [Hasgall, P. A. et al 2015]. The following properties of magnetite were used: magnetic saturation 92.0 kA/m, the anisotropy constant 30 kJ/m3, the surfactant layer thickness 2.0 nm and the density 5180 kg/m3. Log-normal volume weighted particle distribution was assumed (median = 10.3 nm, standard deviation = 0.16). Volume weighted log-normal distribution was used for the MNPs. The spatial distribution of the MNPs in the tumour volume was considered homogenous i.e. the volume of the ferrofluid and the volume of the tumour tissue were the same. The concentration of MNPs in the tumour was raised to 15 mg/ml in order to generate the desired temperature in the tumour area.

The MNP relaxation times were estimated using magnetic field dependent formulas. For Brownian relaxation, the viscosity of blood was used instead of water. When comparing MNPs suspended in solution with MNPs suspended in the tumour tissue, the Neel relaxation time was same. The biological and technical information concerning the rat model can be seen in Table 1. Numerous tissues were added in order to construct a life-like *in vivo* numerical model.

**Table 1:** Biological and technical information concerning the rat model [Christ, A et al 2009].

| Sex | Weight | Length<br>(snout-vent) | Length<br>(total) | Properties | Description | Number<br>of slices | Slice<br>separation for<br>discretization |
| --- | --- | --- | --- | --- | --- | --- | --- |
| female | 502.5 g | 22.5 cm | 38.5 cm | 1 tumor | Sprague<br>Dawley | 738 | 1.052 mm |

## Results and discussion

### Non-contact Temperature Monitor

A schematic can be found in fig. 1b to understand positioning of the probes in this section dedicated to non-contact temperature measurement. Figure 2a plots the temperature variation within the exposure time of a magnetic field of strength 5 mT at an AC frequency of 218.7 kHz, recorded simultaneously using a fibre optic temperature sensor in contact with the radio frequency susceptible nanoparticles and an infra-red temperature sensor connected through the base of a petri dish. Very small differences in the measurements of the absolute temperatures using these two probes indicate that the mid infrared region can measure the temperature of the MNP sample through/ on the permeable material during an AMF exposure. The comparison between the two fibre optic temperature sensors used for recording the temperature variation in the MNPs through the exposure time of the magnetic field is plotted in fig. 2b.

**Fig. 1.**
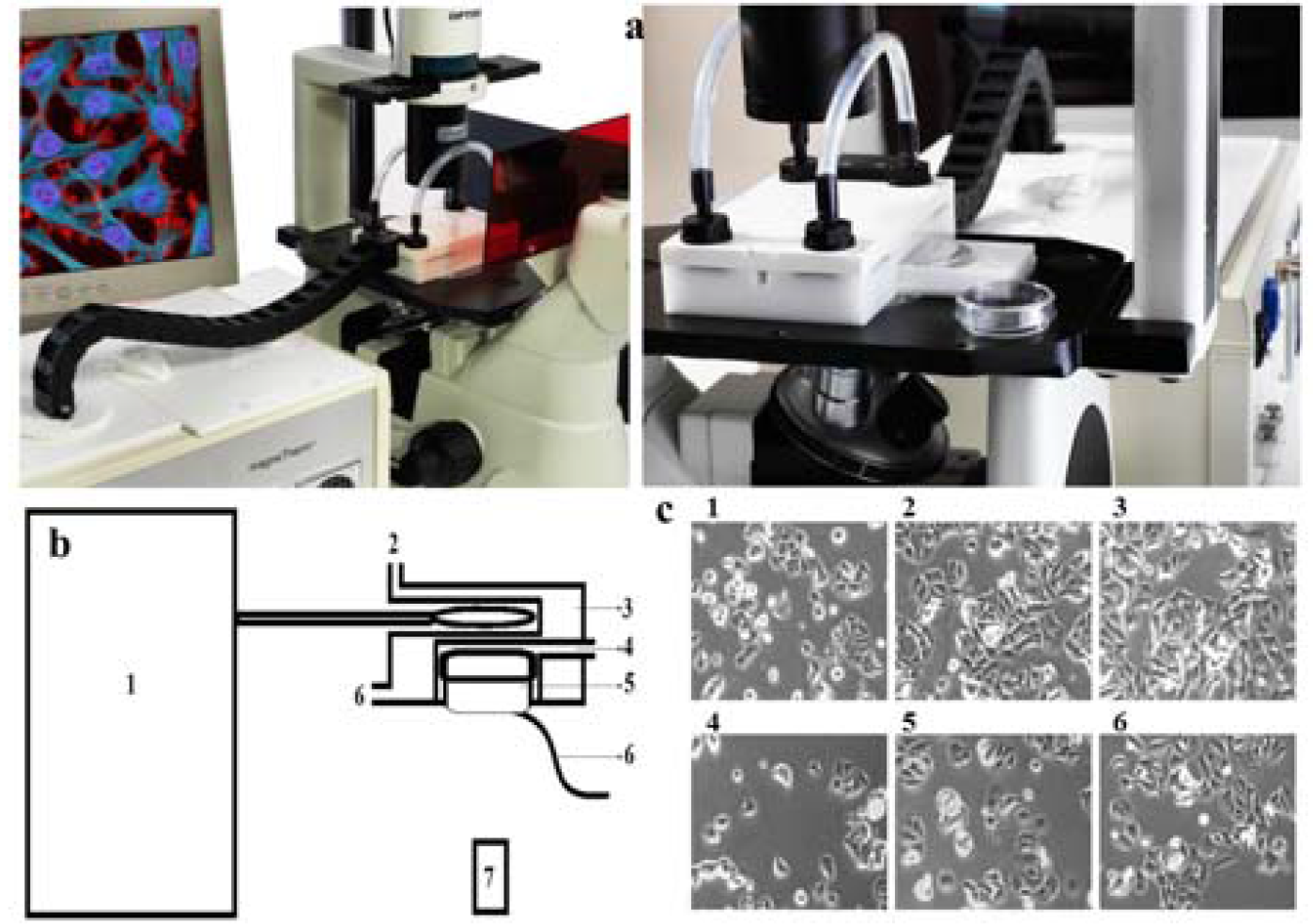
(a) Non-contact temperature measurement setup – the coil system used for real-time magnetic fluid hyperthermia microscopy studies. (b) Block diagram schematic of the proposed technique and the apparatus used. b1 – alternating magnetic field applicator, which can generate time-varying magnetic fields of 1 kHz to 1.2 MHz and 20 mT flux density; b2 – cell-culture incubation chamber’s water inlet; b3 – cell-culture incubation chamber’s water jacket to maintain physiological temperature; b4 – biological gas flow; b5 – cell-culture vessel with 50 μm or less base thickness; further referred to as a 35 mm2 cell-culture-treated petri dish. b6 – cell-culture-incubation chamber’s water outlet; b7 – fibre optic temperature sensor; b8 – infra-red temperature sensor; further referred to as CSmicro 2W Mid-Infrared (MIR) 8-14 μm temperature sensor (Optris, Berlin, Germany). (c) MCF 7 breast cancer cells were seeded on Day 0 and maintained within the setup shown in Fig. 1(a) up to Day 3; c1, 2, 3 – Day 1 to 3 cells grown within the coil setup incubator; c4, 5, 6 – Day 1 to 3 control cells grown in laboratory cell incubator.

**Fig. 2.**
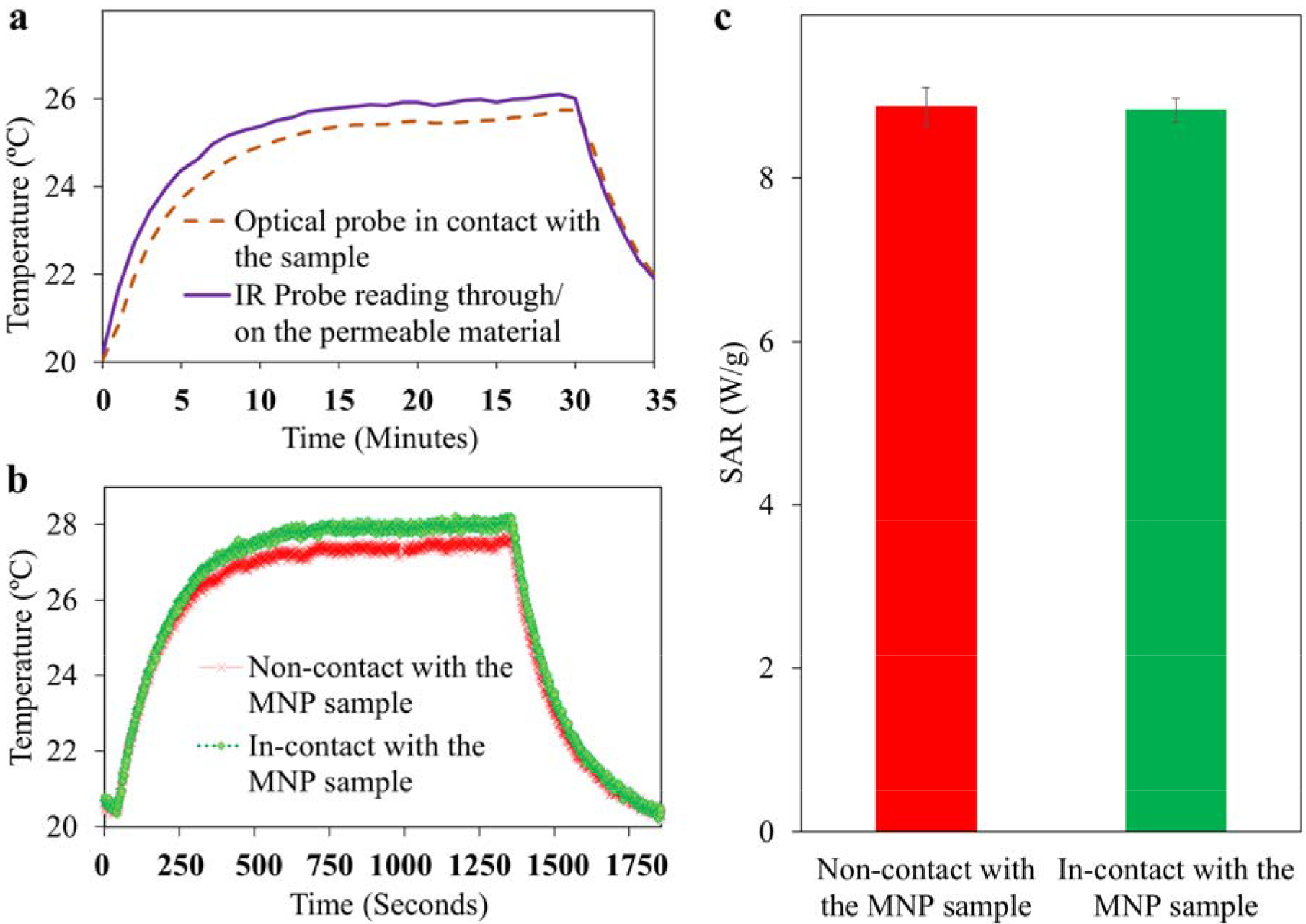
(a) Temperature variation through the exposure time of a magnetic field with strength 5 mT at an AC frequency of 218.7 kHz, recorded simultaneously using a fibre optic temperature sensor (OTG-M360-W-62ST-1.5PTFE-XN-WGT-M2; OPSENS, Quebec, Canada) in contact with the radio frequency susceptible nanoparticles and an infra-red temperature sensor (CSmicro 2WMid-Infrared (MIR), Optris, Berlin, Germany) with a working range of 8-14 μm, connected through the base of the petri dish, (b) A comparison between two fibre optic temperature sensors (OTG-M360-10-62ST-1.5PTFE-XN-10GT-M2, OPSENS, Quebec, Canada) used for recording the temperature variation throughout the exposure time of the magnetic field. One sensing probe was placed in contact with the radio frequency susceptible MNP, and the other measured the temperature through the base of the petri dish without any direct contact with the radio frequency susceptible nanoparticle sample, (c) Specific absorption rate (SAR) values calculated from the changes in temperature with time, using both of the fibre optic temperature sensors (two OTG-M360-10-62ST-1.5PTFE-XN-10GT-M2; OPSENS, Quebec, Canada), one in contact with the radiofrequency susceptible nanoparticle sample and the other connected to the base of the petri dish. There is no significant difference (% 0.5) between the SAR values calculated using the temperatures yielded by the probe in contact and the probe not directly in contact with the sample.

Here, one sensing probe was directly in contact with the radio frequency susceptible MNP, and the other was not in direct contact, but measured the temperature through the base of the petri dish. The temperatures measured by both the in-contact and the non-contact fibre optic probes were used to calculate SAR values for the MNP sample – by considering the entire region of the change of temperature over the entire AMF exposure time using the corrected slope method [Wildeboer, R. R et al., 2014]; this is represented in fig. 2c. The value of SAR which was based on the non-contact temperature measurement, SAR = 8.88 ± 0.24 W/g, agreed well with the value of 8.83 ± 0.15 W/g, as calculated using the temperature measurement probe which was in contact with the MNP sample. The method being developed here, demonstrated its ability to accurately perform non-contact temperature measurement. It is a suitably cost-effective method for the non-contact monitoring of *in vitro* samples and could make redundant the complex algorithms that have been proposed [Silva, L. R. and Stuart, M. D 2016] to enable the use of non-contact temperature measurement via mid-infra-red pyrometers – these cannot measure temperature through the commonly used polypropylene sample tubes and glass petri dishes. The advantage of this type of setup is the ability to perform non-contact temperature measurements using a fibre-optic sensor, an infra-red pyrometer and an infra-red camera. However, the petri dish itself is a limitation of this experimental setup as different types of MNP samples have varying heat dissipation profiles and the thin polystyrene (or other polymer) layer – 70 °C will have a temperature-resistant tolerance. Furthermore, near infra-red wavelength window of the pyrometer deployed here only records surface temperature.

### Drug Release System

The whole setup was designed using CAD and fabricated to fit within the solenoid coil setup as commercially available components did not fit the required specifications. Fig. 3a highlights the unique features of the proposed drug release testing setup including: (i) a disposable dialysis tube holding a shaft capable of accommodating a 250 μL – 3 mL volume of sample; (ii) a high resolution fibre optic temperature sensor for real time temperature measurements; (iii) the sample-holding shaft driven by a stepper motor to allow continuous stirring of the solution – for enabling even distribution of the drug released into the dissolute; (iv) the sampling port/shaft for manual spectrophotometer analysis and to accommodate a custom engineered non-metal fibre optic probe for real time UV-VIS spectrophotometer analysis, while performing the AMF mediated drug release experiment; and (v) a water jacket enclosing the dissolute holding chamber to maintain ambient temperature in the dissolute as well as in the MNP + Drug sample. These features are appropriate for the design of a working prototype (as shown in fig. 3b-d) and for recording SAR measurements at high precision.

**Fig. 3.**
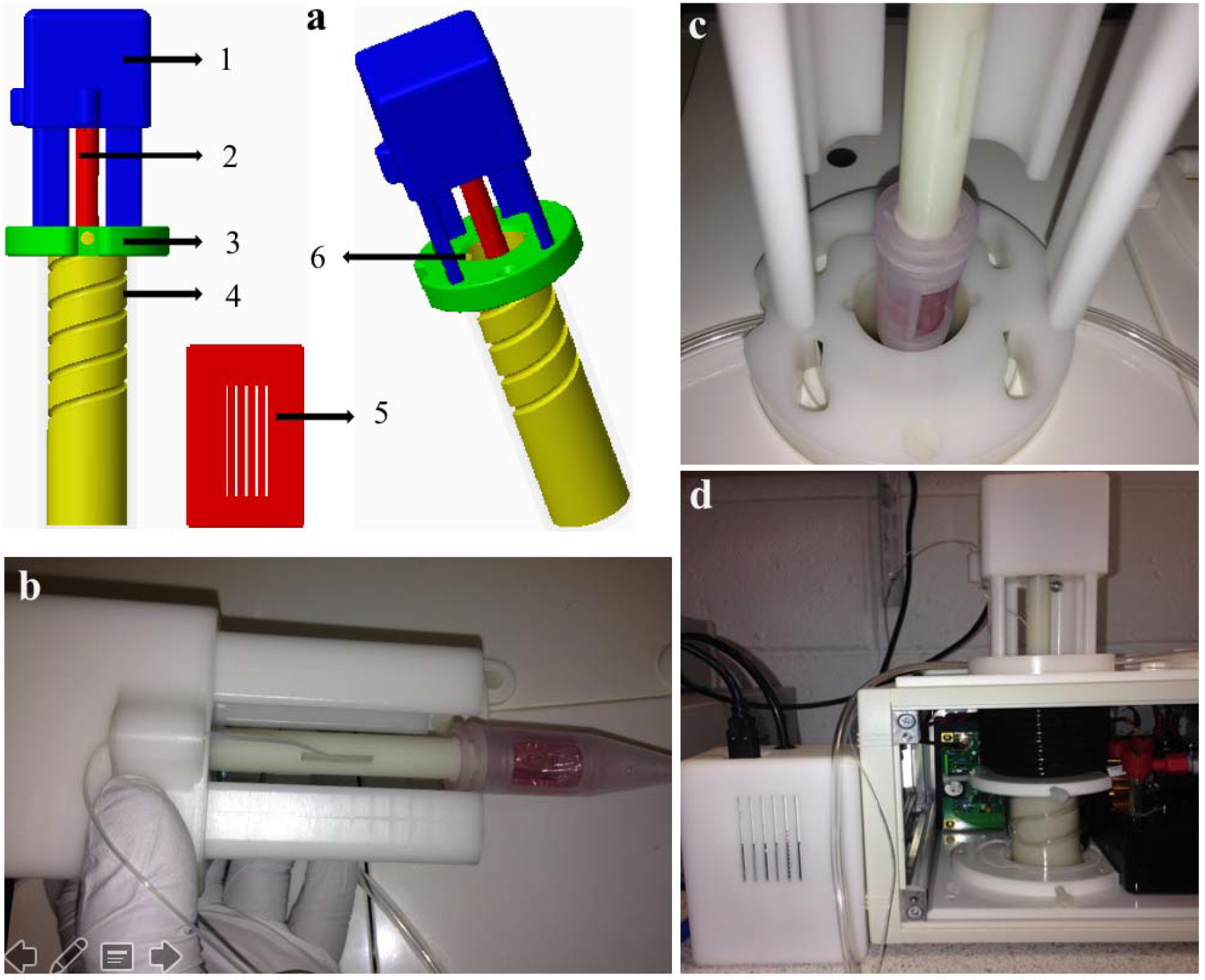
Various parts of the AMF mediated drug release testing setup (a): (a1) the stepper motor mounting; (a2) the dialysis tube sealing shaft with an aperture for a temperature probe attached to the stepper motor, (a3) a magneTherm^™^ mounting clamp; (a4) the water jacket (bottom sealed); (a5) the control box; and (a6) the sampling port. (b) A photograph of the AMF mediated drug release analysis setup the stepper motor attached to the 12 kDa dialysis tube through a shaft – the shaft also provides temperature probe access to the MNP sample; (c) the dialysis tube attached to the stepper motor positioned within the dissolute; (d) and the complete setup with the water jacket attached to the magneTherm^™^ coil enclosure including the microprocessor (within control box), programmed to provide oscillating motion.

Here, the steps produced by the stepper motor are critical, as an increased rotation per minute (rpm) rate might increase heat dissipation by the sample into the surrounding dissolute. Additionally, the volume of dissolute is critical as the stimulus involved for drug release is heat. Generally, 20 rpm and 25 ml of dissolute volume were used in this study. This dissolute volume was used so that the dialysis tube stayed immersed [Hervault, A. et al 2016], and is 12.5 times more than the MNP loaded with doxorubicin (MNP + Dox) sample volume. Fig. 4a shows the recorded temperature over time curve for 50 nm sized maghemite nanoparticles (2 ml of 20 mg/ml) dispersed in 100 μl doxorubicin. The drug release testing setup facilitated heating of the MNP containing solution from 23.91 °C to 67.43 °C after exposure to AMF of strength 16 mT with frequency 107.7 kHz, for 60 minutes, even with the dialysis tube immersed in dissolute. A proof of principle experiment shown in fig. 4b demonstrates the usability of this setup which has been designed for alternating magnetic field mediated drug release studies.

**Fig. 4.**
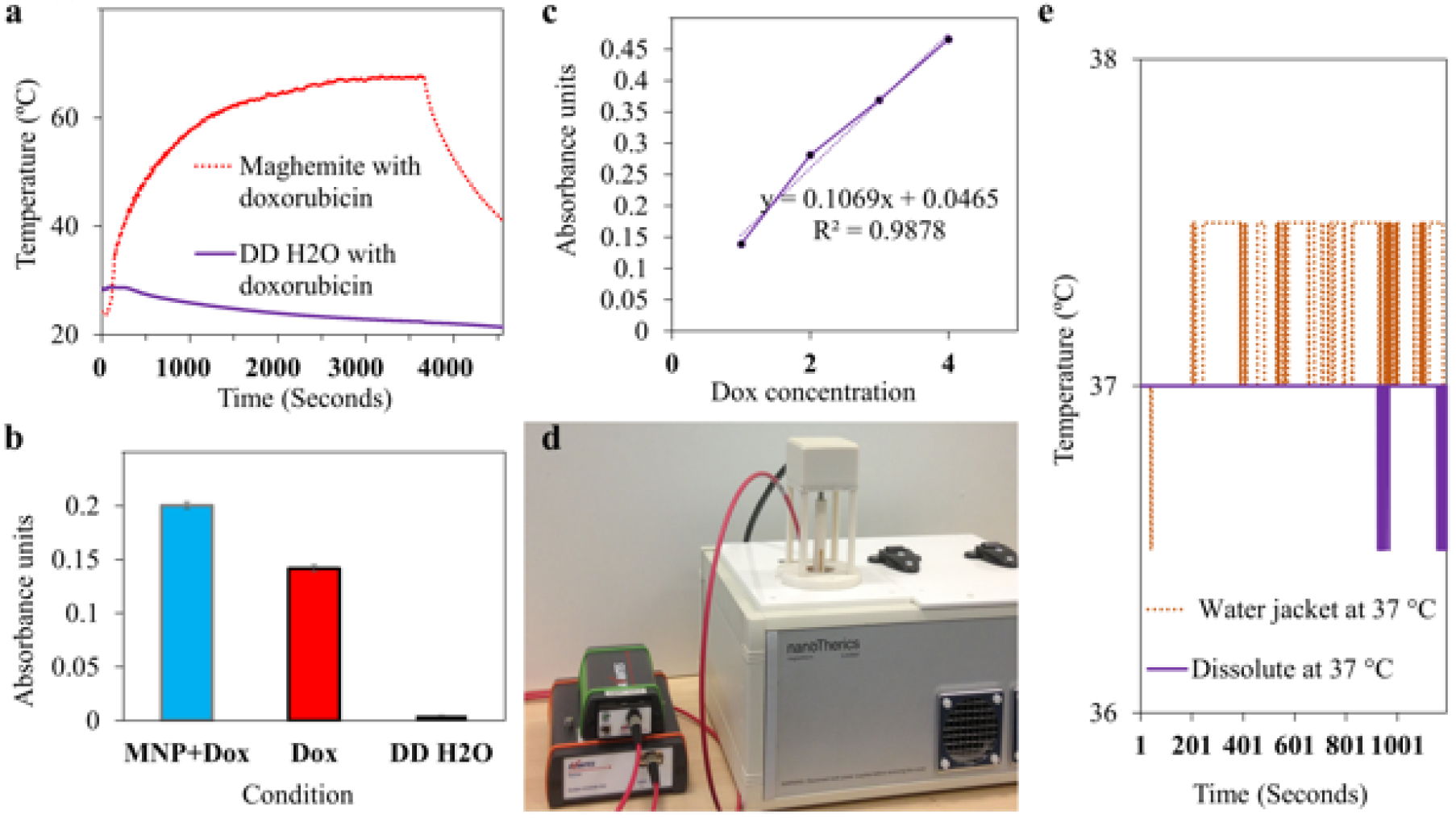
(a) Real time temperature changes in 50 nm sized maghemite nanoparticles (2 ml of 20 mg/ml) dispersed in 100 μl doxorubicin via a dialysis tube containing 25 ml of double de-ionized water (DD H_2_O); this was recorded using the AMF mediated drug release analysis setup. A fibre optic temperature sensor was used for real time temperature measurement and the sample holding shaft was set to 20 rotations per minute. (b) Dox release measurement after 1 hour of exposure to AMF at strength 16 mT and frequency 107.7 kHz followed by cooling for 15 minutes by maintaining the stirring rotation time at 20 rpm – the real-time temperature measurement was recorded using an Osensa optical sensor. (c) Calibration curve for varying concentrations of doxorubicin. (d) The camera view of the experimental setup used for testing the AMF mediated drug release experiment: connected to a light source, a real time UV-Vis absorption spectroscopy probe (compatible for AMF), and a spectrophotometer interface. (e) Temperature measurements for the water jacket without the dissolute and with the dissolute enclosed within the water jacket used in the drug release testing setup.

The results show that the heat generated using the maghemite magnetic nanoparticles exposed to an alternating magnetic field has enhanced doxorubicin – cancer drug release *in vitro* (0.200 abs units) when compared to a doxorubicin-only control (0.142 abs units). The real-time temperature data (37 °C baseline temperature) from fig. 4e shows that it is possible to maintain a desired ambient temperature using the water jacket module which is utilised with the drug release testing setup. The rotation can be adjusted from 120 to 20 RPM to mimic in vivo blood flow. The design suitability was further enhanced by a water jacket fabricated in clear material to enable visualization of the released drug into the medium. Displacement spacers were added to the water jacket design to perform experiments with different volumes of drug release medium.

The steps produced by the stepper motor and volume of dissolute are critical factors. Variation of volume of dissolute does influence the heat dissipation from the MNP + Drug conjugate. Moreover, the heat dissipation from the MNP: Volume of dissolute ratio depends upon the magnetic and chemical properties of the MNP + Drug conjugate and the flux parameters of the magnetic field applicator. Herein, aqueous suspension of commercially available 50 nm sized maghemite particles were conjugated with Doxorubicin and exposed to AMF to demonstrate the function of the proposed design in vitro. The change of temperature curve shows that heating ranges from 23 to 67 °C after AMF exposure for this MNP + drug conjugate. The temperature range was wide and hence not practical for in vivo applications, as the temperature was constantly increasing due to continuous AMF exposure over an hour. However, the properties and experimental settings will vary for custom-engineered MNP and in vivo studies. Nevertheless, fig. 4 demonstrates the utility of this design for AMF-mediated drug-release studies.

While solenoid coil – AMF systems have been used for heating Doxorubicin-coated MNPs to demonstrate the heating properties of MNPs [Benyettou, F et al 2016], recent studies suggest that such solenoid coil setups will be suitable for AMF-mediated controlled drug-release studies [Mertz, D et al 2017]. Moreover, the authors of this recent review propose a theoretical model for carrying out drug release experiment through drug diffusion across semipermeable membrane and this model correlates with the drug release analysis setup design discussed in our study. However, they suggest the following: The setup used for in vitro AMF mediated drug release should involve a high-power RF-AC magnetic field system with a coil, whose diameter can accommodate 1-5 ml of MNP-drug conjugate. The MNP-drug conjugate enclosed within a dialysis bag should be surrounded by release medium with volume at least 10 times that of the sample. The molecular weight cut-off of the dialysis membrane has to be high but should be able to withhold the MNP-drug conjugate, as the drug-release kinetics are limited by the permeability of the surface-functionalized Nano carriers. Use of a non-metallic temperature probe, a water jacket for maintaining physiological temperature, and drug-release media circulation to mimic blood circulation, in addition to caution for eddy current mediated non-specific heating of the sample, was advised. Furthermore, they added that monitoring the drug release profile continuously, while keeping the AMF ON will be beneficial for post experiment data analysis, as then the data will be derived from aliquots at specific intervals [Mertz, D et al 2017]. Moreover, all of the aforementioned points were incorporated into the design of the AMF-mediated drug-release system discussed in this study.

### Double-stacked Planar Coil System Design

The numerical calculations (as plotted in fig. 5a-f) for the 2-layers stacked coil with 14 turns, result in an estimate for absolute flux density of |B| = 51 mT and for RMS flux density of B_rms_ = 36 mT, at coil current of 56 I_rms_. This design demonstrates a potential of achieving higher flux densities with a relatively low amount of current through the coil and a reduction of heat generated by Joule’s heating while current flows in the coil, when compared to the commonly used water-cooled induction coils (solenoid and planar). The highly intense and homogeneous flux line patterns at the centre of the coil (at the sample position) suggest that this is an ideal coil design for calorimetry experiments. The numerical simulations of 2 × 16 turn and 2 × 15 turn coils did provide better flux density, homogeneity, and intensity in the centre of the coil, as compared to those of a coil with 14 turns. Calculations based on 2 × 15 coils yielded an absolute flux density of |B| = 54.3 mT and an RMS flux density of B_rms_ = 38.4 mT; and calculations based on 2 × 16 coils yielded an absolute flux density of |B| = 57.9 mT and an RMS flux density of Brms = 46.3 mT; all these values were obtained for a coil current of 56 I_rms_.

**Fig. 5:**
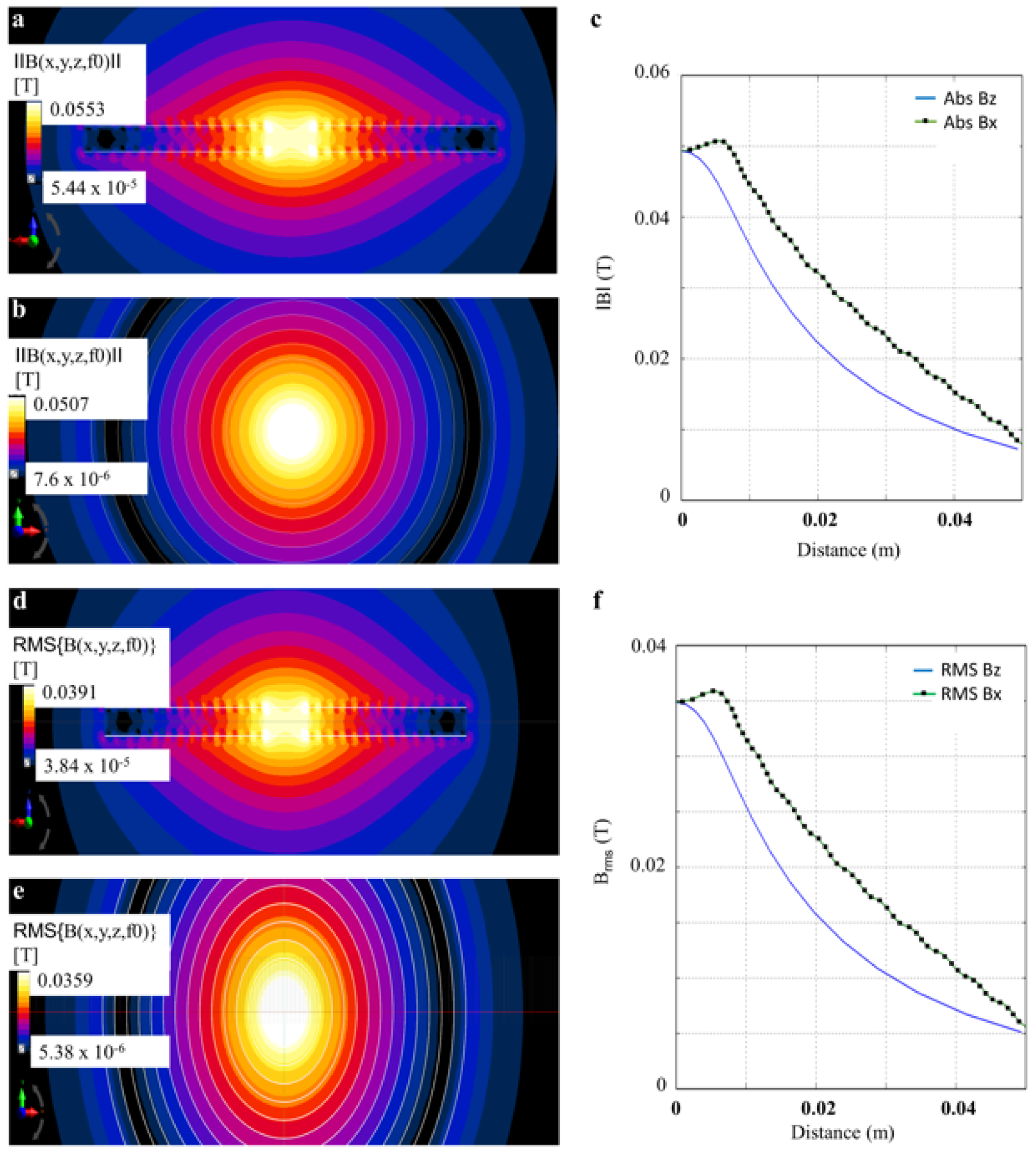
(a) The simulated contour plots for the double stacked planar coil with 14 turns in each stack for: the xy plane absolute flux density (|B_xy_|), (b) the zy plane absolute flux density (|B_Zy_|), and (c) the absolute flux density in the xz plane of the coil applied with I_rms_= 56 A, |B| = 51 mT. (d) The xy plane RMS flux density (B_xy_), (e) the zy plane RMS flux density (B_zy_), and (f) the RMS flux density in the xz plane for the coil applied with I_rms_ = 56 A, B_rms_ = 36 mT.

**Fig. 6.**
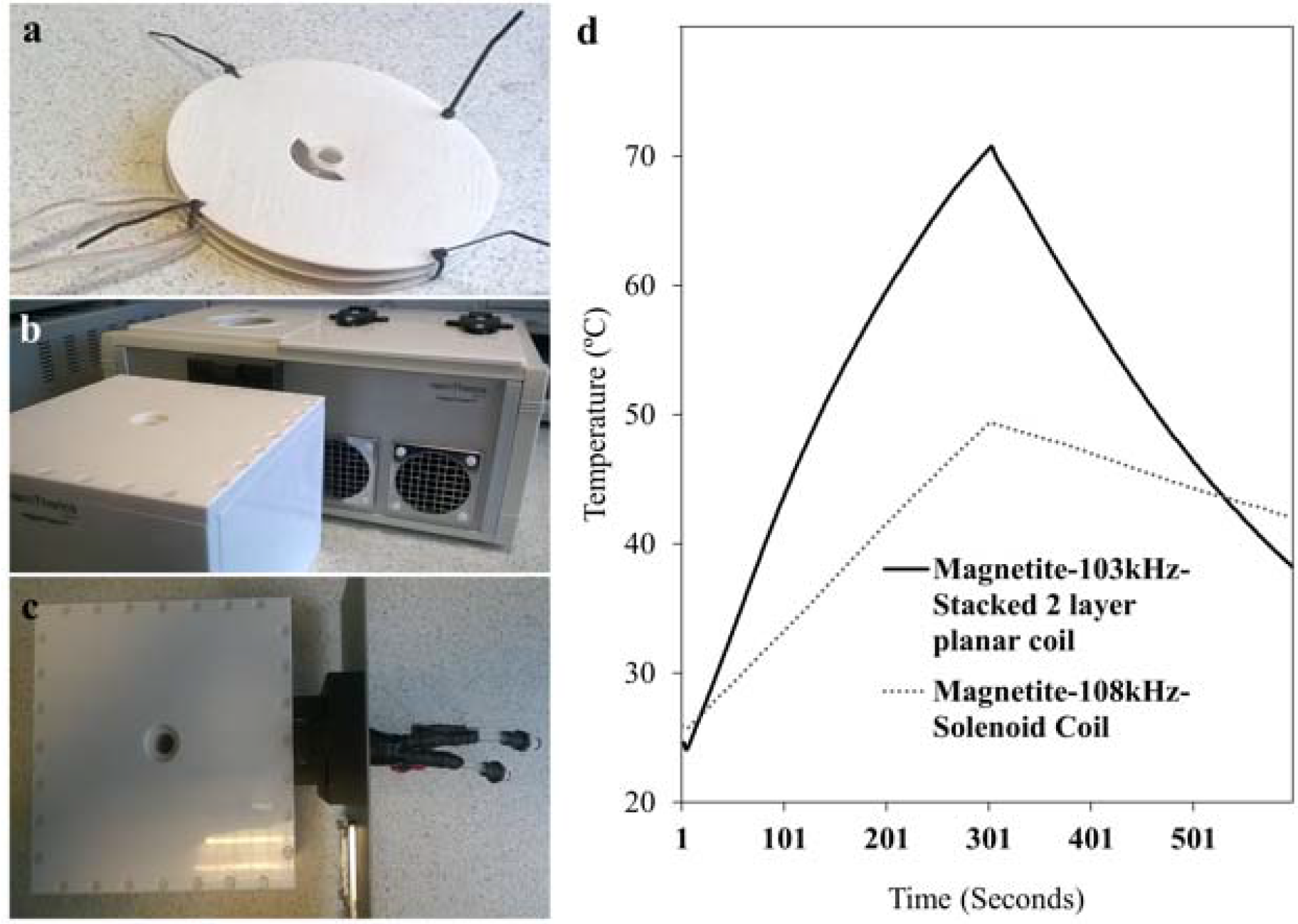
(a) The fabricated high flux density coil setup: the stacked 2-layer planar coil; (b) the coil mount, to hold the stacked 2-layer planar coil, using a sample aperture; (c) and the Set up mounted externally onto the magneTherm^™^ system, (d) Change of temperature over time graph for a 5 mg /ml aqueous suspension of DMSA stabilised 10.3 nm sized magnetite nanoparticles. A calorimetric experiment comparing double stacked planar coil with a commercial setup using a simple solenoid at 56 I_rms_ coil current.

However, because of fabrication issues, we used a 2 × 14 turn coil design (as shown in fig. 6a-c) without compromising any measurements. A solenoid coil has usually been considered the appropriate choice in commercial instruments marketed for this purpose [Macías-Martínez et al. 2016; Iacob, N. et al 2016; Calatayud, M. P et al 2016; Soares, P. I., et al 2016]. Hence, this high field-strength module was compared with a standard solenoid coil – the magneTherm system with dimensions D_t_ = 4.06 mm, d_t_ = 2.68 mm, N = 17, l = 2, S_N_ = 1.04 mm, S_1_ = 2.08 mm, d_c_ = 51.66 mm, D_c_ = 73.62 mm, and H = 49.44. This solenoid coil, however, produced a RMS B flux density of about 16.71 mT at 56 I_rms_ coil current. The stacked planar coil did heat the MNP sample much more efficiently than did the solenoid coil – as seen in fig. 6d. The HFS coil design discussed here can be used for calorimetry as the centre of the coil can accommodate sample sizes up to 2 ml. Though the design does not include an adiabatic shield [Natividad, E et al., 2011; Natividad, E et al., 2009], either vacuum layer or foam insulation can be included between the dielectric wall and the sample tube if the sample tube size is reduced to accommodate 1 ml from the 2 ml used in the study. However, it is evident from the fig. 5; 6d, that this design provides enhanced effects in terms of incident flux intensity, homogeneity and flux density for the same coil current, when compared with a solenoid coil (with wide sample aperture) [Subramanian, M 2016]. These parameters are the reason for observing increased heat dissipation in the calorimetry comparison results. In addition to induction coils, magnetic core-mediated heating is also plausible for MNP calorimetry experiments [Tai, C.C. and Chen, C.C., 2008]. It is also possible to use a flux concentrator to influence the field parameters [Ivkov, R et al., 2005].

### Curved planar coil - in vivo set up

In MFH, a low cost thermal imaging technique has been successfully used for obtaining real time in vitro and in vivo surface temperature measurements [Mannucci S et al., 2014; Corato R D et al., 2015; Espinosa A et al., 2016]. This is beneficial for heat dosimetry and clinical planning. It is possible to estimate the power dissipation from MNPs and intratumoral temperature increase based on the surface temperature measurements [Rodrigues HF et al 2017]. Furthermore, the technique is non-contact and covers a wide area, which is a major advantage over point-measurement techniques [Lahiri BB et al., 2016]. Thus, an efficient applicator design that will enable performance of in vivo AMF exposure and thermal imaging simultaneously is required.

This in vivo coil design was inspired by the shim coil used to adjust homogeneity of a static magnetic field by changing the current flowing through it for high resolution MRI. The curvature of the coil will depend upon the in vivo water jacket OD, and the geometry of the in vivo coil used for calculations is shown in fig. 7a. The curved planar coil *in vivo* set up (the CAD model) and the flux density calculations can be seen in fig. 7. Calculations revealed that the incident flux density is non-homogenous and failed to cover the whole of the animal model (Fig. 7d). The flux density in the z axis ranged up to μ_0_H = 31.42 mT (Fig. 7c).

**Fig. 7:**
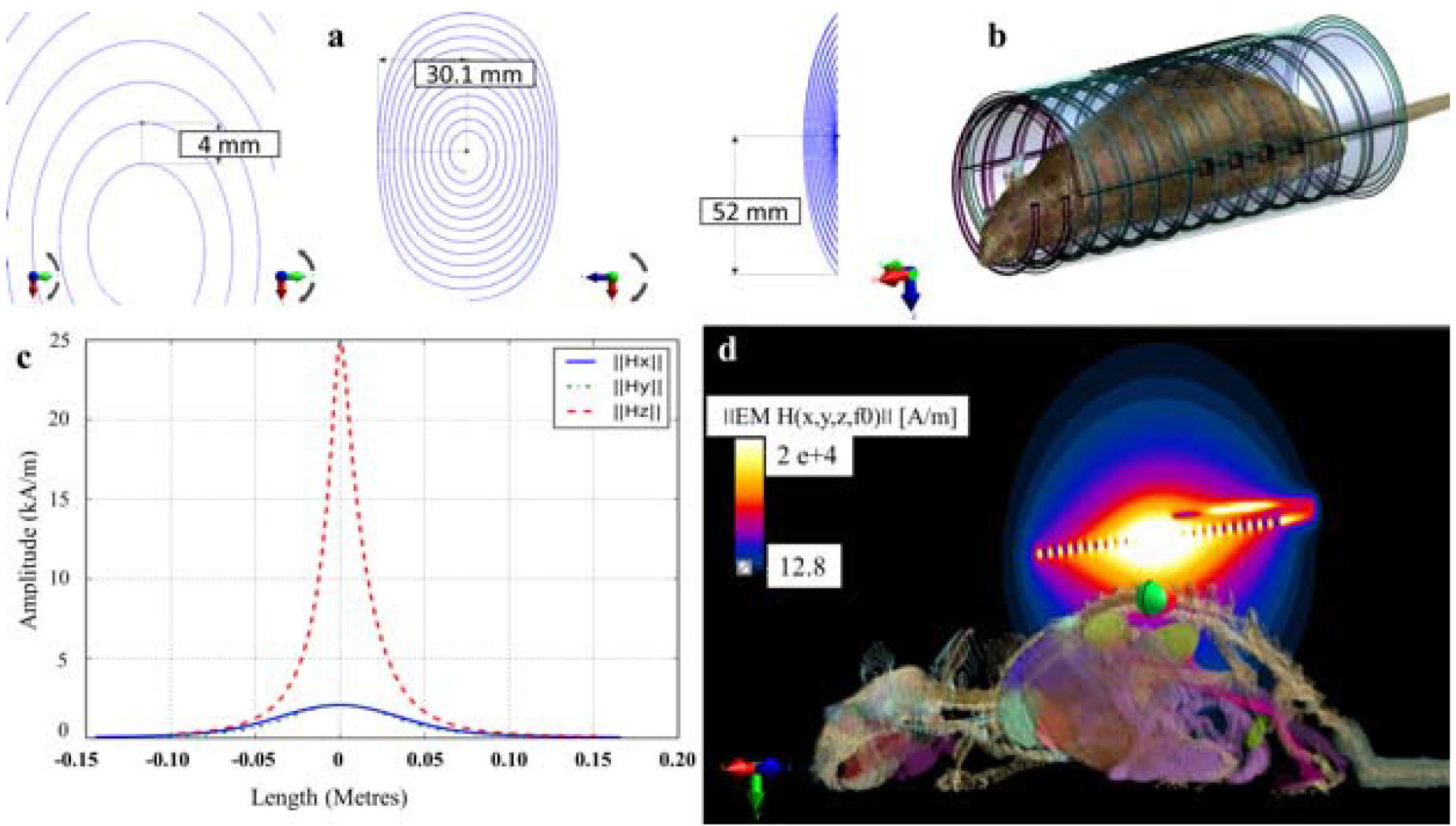
The curved planar coil setup for the in vivo experiments: the flux density numerical calculations regarding the setup for 56 Amps (RMS); (a) geometry of the curved planar coil (b) an anatomically correct 3D rat model exposed to AMF generated by the curved planar coil setup with a coil current of 56 Amps (RMS) together with the viewing window for thermal imaging and the coil-shaped water jacket for maintaining physiological temperature can be seen in the in vivo coil setup - CAD model; (c) Magnetic field strength (H) along the main axis with regard to the coil middle; (d) Magnetic field strength XY-cross section passing through the middle of the coil (green ellipsoidal shape indicates the tumour position).

Magnetic hysteresis modelling is complex and Linear Response Theory (LRT) for correlation with experimental data has been under scrutiny, since characterization of bulk and microscopic properties (static and dynamic), such as intercore dipole interactions and differences in internal magnetic (intracore) structure, will affect the heating performance of the ferrofluids. Measurements limited to amplitudes lower than saturation usually fail to establish the nonlinearity of power loss [Dennis CL et al., 2015] and stirring-mediated power loss due to aggregated nanoparticles in an aqueous suspension cannot be neglected [Vallejo-Fernandez G et al., 2013]. However, in recent years, LRT has been evaluated and modified for magnetic hysteresis modelling considering the importance of the dispersity index and the nanoparticle anisotropy constant [Boskovic M et al., 2015]. Furthermore, high nanoparticle concentrations were found to correlate with increasing chain length and when magnetic dipolar contribution was considered, a decrease in hyperthermia efficiency was demonstrated [Branquinho LC et al., 2013]. Moreover, the heat dissipation due to susceptibility loss is low and can be neglected, with hysteresis loss being the dominant mechanism based on the power loss measurements and calculations performed on the HyperMAG^®^ particles [Vallejo-Fernandez G et al., 2013] used in this study. Nonetheless, discrepancies with estimated parameters and theoretical predictions arising due to particle-particle interactions require normalization. In addition, it must be acknowledged that accurate prediction of SAR is difficult [Verde, E.L et al., 2012; Verde, E.L et al., 2012; Ruta S et al., 2015].

The physical evaluation of the prototype using a high frequency magnetic field probe did correlate with the magnetic field calculations. Furthermore, as an example application, the magnetic field strength on the tumour surface was calculated as it is shown in Fig. 7d. This magnetic field can be considered as the effective one which leads to further phenomenon like the eddy currents (see Fig. 8b), the magnetic power dissipation (see Fig. 8c) and finally to the temperature distribution within the tumour tissues (fig. 8d). One should notice that the aforementioned phenomenon take place in the whole rat model and can be considered as the whole-body hyperthermia. Only the magnetic power losses can be controlled when magnetic nanoparticles are injected into the tumour in order to emphasis the temperature rise in the target area.

**Fig. 8:**
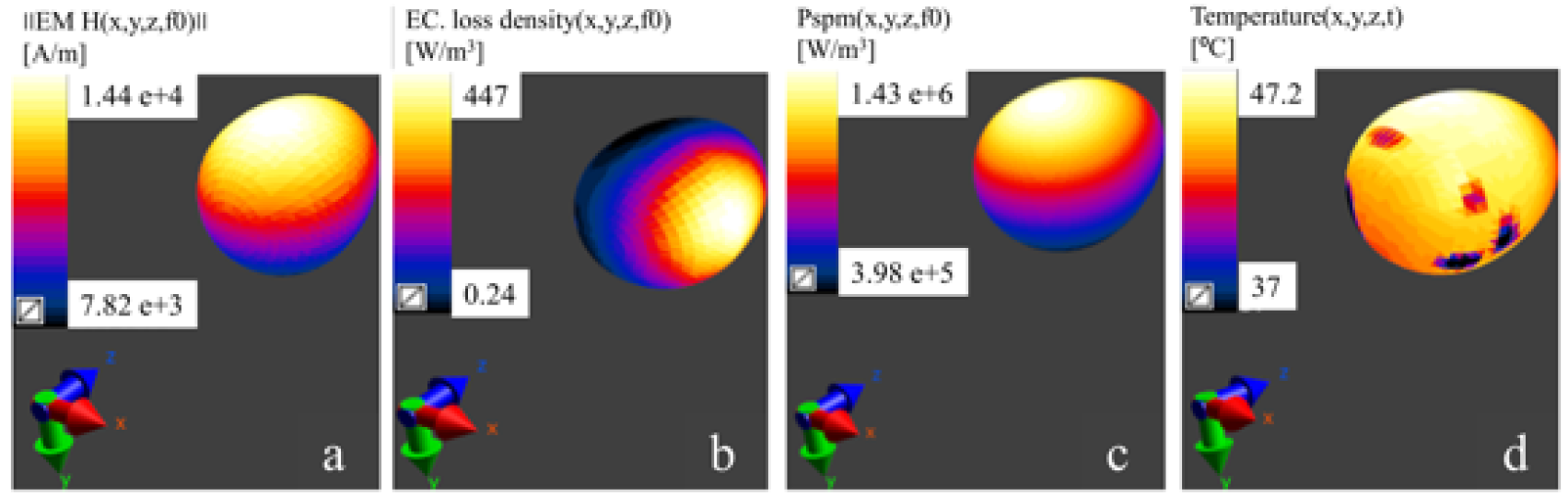
Numerical calculations relative to the tumour from Fig. 7d: (a) Magnetic field strength on the tumour surface; (b) power density due to the eddy currents; (c) magnetic power loss density and (d) temperature distribution on the tumour surface after 1000 seconds of exposure.

In the case of presented setup, which was designed for *in vivo* experiments, it was able to heat a 5 mg /ml aqueous suspension of DMSA stabilised 10.3 nm sized magnetite nanoparticles by 3 - 5° C within 5 minutes, when a 2 ml sample containing tube was placed in proximity to the centre of the coil. However, on the base of computer modelling it was found that 5 mg/ml of magnetic nanoparticles concentration was too small to rise the temperature within the tumour. That is why the concentration was increased to 15 mg/ml leading to magnetic power losses up to 1.4 MW/m^3^ and the temperature rise up to 47 °C in the tumour (see Fig. 8c-d). The following data were used in order to calculate the total magnetic power: the volume weighted log-normal distribution, the magnetic saturation 92.0 kA/m, the anisotropy constant 30 kJ/m^3^, the surfactant layer thickness 2.0 nm and density 5180 kg/m^3^ which are equivalent to the physical properties of the magnetite. Moreover, the tumor tissue with MNPs was treated as a homogeneous composite.

Even though the proposed design would be ideal for real time full body thermal imaging while simultaneously maintaining physiological temperature within the animal chamber, the heating efficiency of the curved planar coil, and the homogeneity of the incident flux density was low when compared to the aforementioned 17 turn solenoid coil (11 ° C within 5 minutes). Hence, further investigation of the coil design is required in order to improve its efficiency for such experiments. Though, the numerical calculations performed in this study will help researchers in planning *in-vivo* experiments using such anatomically correct female-rat models with tumours.

Fig. 8 shows the numerical calculations relative to the rat 3D model placed within the curved planar coil setup. Studies based on nanoparticles with equal saturation magnetization but varying magnetic anisotropy, suggest that the latter parameter plays a key role in the transition from the linear to non-linear regime when the flux density is increased. Calculations based on Linear Response Theory, the Stochastic Landau-Lifshitz method and kinetic Monte-Carlo models have been used to understand the effect of material properties and inter-particle interactions on specific heat power. Such studies help us engineer magnetic fluids with better magneto-thermal properties for hyperthermia applications. However, discrepancies with estimated parameters and theoretical predictions arising due to particle-particle interactions need to be accounted for. And we need to acknowledge that accurate prediction of SAR is difficult [Verde, E.L et al., 2012; Verde, E.L et al., 2012; Ruta S et al., 2015].

Nevertheless, such in silico evaluations will provide knowledge to design suitable AMF applicators for in vivo/ in vitro MNP heating as demonstrated in this study.

## Conclusion

The measuring techniques and instrumentation developed here, demonstrated a cost-effective, accurate means of performing non-contact temperature measurements on *in vitro* samples. Likewise, the proposed drug release analysis setup will allow researchers to perform and replicate their AMF mediated drug release experiments with accuracy and repeatability. We have also shown that the water-cooled stacked planar coil system may be an efficient tool for performing MNP-calorimetric experiments. However, the described *in vivo* coil setup does require further development. Nevertheless, the aforementioned results infer that the tools and instrumentation proposed in this article has the potential to allow standardization of a variety of AMF-mediated MNP experiments with efficiency and repeatability.

## Author contributions

MS conceived the idea, designed and fabricated all the instruments described in the above article, MS, AM, OH performed the evaluation experiments. *In vivo* numerical modelling was designed and done by AM. MS, AM and JD performed data analysis. MS Wrote the manuscript. AKM and JD edited the manuscript.

## Conflicts of interest

The research was funded by nanoTherics Ltd. nanoTherics is involved with commercialisation of live cell - alternating magnetic field (LC-AMF) setup, drug release analysis system and high flux density module. M.S. worked as a research scientist in bioengineering for nanoTherics, and J.D. is a consultant for nanoTherics. Currently, M.S. is funded by the EPSRC and Imperial College London. A.M., A.K.M and O.H are academic researchers and are unrelated to nanoTherics.

